# The acute sleep-inducing effects of light require histamine neurotransmission in mice

**DOI:** 10.1101/2025.06.26.661495

**Authors:** Yanlong Hou, Ni Tang, Sophie Wu, Yufeng Shao, Jian-Sheng Lin, Claude Gronfier

**Author notes:** These authors contributed equally to this work (co-first) (Yanlong Hou, Ni Tang, Sophie Wu). These authors contributed equally to this work (co-last) (Jian-Sheng Lin, Claude Gronfier). Corresponding author **Corresponding Authors: Claude Gronfier**, Lyon Neuroscience Research Center, Waking team, Inserm UMRS 1028, CNRS UMR 5292, Université Claude Bernard Lyon 1, Université de Lyon, F-69000, Lyon, France,; **Jian-Sheng Lin**, Lyon Neuroscience Research Center, Waking team, Inserm UMRS 1028, CNRS UMR 5292, Université Claude Bernard Lyon 1, Université de Lyon, F-69000, Lyon, France.

## Abstract

Sleep regulation depends on the complex interplay between homeostatic and circadian processes synchronized by the light/dark cycle. Sleep is also directly regulated by light via the retinal inputs to the preoptic area (POA). Although the light-responsive POA neurons project to several wake-promoting structures, including histaminergic neurons in the tuberomammillary nucleus (TMn), there is no functional evidence for their involvement in light-induced sleep. To bridge this gap, we used histidine decarboxylase (HDC, the histamine-synthetizing enzyme) knockout mice (HDC−/−, n=7) and hM4Di-HDC-cre mice (HDC+/+, n=8) subjected to an ultradian light/dark protocol (LD 1h:1h over 24h), and another group of hM4Di-HDC-cre mice (n=8) exposed to a 1-h light pulse. We found that light pulses during the biological night enhanced slow wave sleep and increased cortical EEG power in the delta range (0.5-3Hz), and that these effects were significantly attenuated both in HDC−/− (83 vs 23 min/6h, p=0.005) under LD 1h:1h condition and in hM4Di-HDC-cre mice after acute chemogenetic silencing of histamine neurons by the DREADD ligand deschloroclozapine (15 vs 6 min/h, p=0.0016) under a 1-h light pulse. In addition, the sleep-inducing effect of light was circadian dependent, with the strongest effect at the beginning and end of the night but no effect at all during the biological day in HDC+/+mice. Our study provides functional evidence that the acute sleep-inducing effects of light on sleep require histamine neurotransmission in mice.

## Introduction

Sleep regulation is driven by two major processes: process C, emanating from the circadian system and process S, sleep homeostasis (Borbély, 1982). The circadian process, driven by a master clock located in the suprachiasmatic nucleus (SCN) of the hypothalamus, orchestrates the timing of sleep and wakefulness in alignment with the natural 24-hour light-dark cycle (Turek, 1985). Concurrently, the homeostatic process builds upon the accumulation of sleep pressure during wakefulness, which is dissipated during sleep, ensuring a balance between sleep and wakefulness (Franken & Dijk, 2024)

Emerging evidence suggests that the regulation of sleep and wakefulness is not solely governed by internal processes but is also impacted by external factors, such as light exposure (Borbély et al., 1975; Dijk et al., 1991; Cajochen et al., 2005; Tsai et al., 2009; Prayag, Jost, et al., 2019; Prayag, Münch, et al., 2019; Prayag, Najjar, et al., 2019; Hubbard et al., 2021). Light is therefore not seen any more as only the most powerful synchronizer of the circadian system in mammals, but also as a powerful sleep regulator. Mechanistically, intrinsically photosensitive retinal ganglion cells (ipRGCs) are essential for detecting environmental light and transmitting the photic information from the retina to the SCN, synchronizing the endogenous clock with the external light-dark cycle, but also activating a wide range of brain structures involved in sleep and alertness, memory, cognition, heart rate, melatonin release and body temperature (Cajochen et al., 2005; Rüger et al., 2006; Chellappa et al., 2011; Hu et al., 2022). The sleep-inducing effect of light require both classical (rods, cones) photoreceptors and ipRGCs (Altimus et al., 2008; Tsai et al., 2009) and its magnitude is circadian-dependent (Tsai et al., 2009).

A recent study found that activation of the POA projecting ipRGCs or of the light-responsive POA neurons both increased slow-wave sleep, disclosing that the ipRGCs-POA circuit mediates the acute effects of light on sleep promotion (Zhang et al., 2021). The sleep-promoting POA must therefore be considered as another component of the neural networks by which light induces sleep in nocturnal species. Besides, these light-responsive POA neurons send inhibitory and monosynaptic inputs to wake-promoting regions such as the lateral hypothalamus (LH) and tuberomammillary nucleus (TMn) (Reitz & Kelz, 2021). However, there is no evidence hitherto that these wake-promoting regions and neurons are functionally involved in the acute light effect on sleep.

Histamine (HA) neurons in the TMn of the posterior hypothalamus promote wakefulness via their widespread projections to the whole brain. They fire tonically and specifically during waking and remain silent during sleep (Vanni-Mercier et al., 2003; Takahashi et al., 2006). Impairment of HA transmission is associated with sleep disorders such as narcolepsy and sleepiness in humans and animals (Haas et al., 2008; Parmentier et al., 2016; Franco et al., 2019). However, whether HA neurons are involved in the sleep-inducing effects of light is unknown.

In this study, we took advantage of a mutant mouse line (histidine decarboxylase, the histamine-synthetizing enzyme, HDC −/−) to investigate whether long-term loss of HA affects the sleep-inducing effect of light, and whether the effect is circadian dependent. We also established a hM4Di-HDC-cre mouse model coupled with a novel DREADD ligand Deschloroclozapine (DCZ, Nagai et al., 2020) to assess the acute effects on sleep of selective silencing HA neurons. Our goal was to test the hypothesis that light induces sleep by inhibiting the HA wake-promoting system, and not only by activating the POA sleep-promoting systems (Zhang et al., 2021). Overall, we tested whether and how the brain neural circuit of sleep induction by light functionally involves HA wake-promoting neurons, as part of the proposed retinal ipRGCs-POA sleep-promoting pathway (Zhang et al., 2021).

## Results

### Histamine mediates the effect of light on sleep and EEG

To investigate the sleep-wake effects of light in both mouse strains and to access the involvement of HA transmission, we exposed mice to the LD1h:1h alternating over 24 h. The impact of light was determined by hourly analysis and by aggregating the total amounts of each sleep stage during the six 1-h light pulses and the six 1-h dark pulses, respectively, throughout the biological day and night and over the 24h (**Fig. 1**). We found that both genotypes showed a clear circadian profile of sleep and wakefulness (W), with more sleep during the biological day (a.k.a. subjective day) than during the biological night (a.k.a. subjective night), as expected for nocturnal animals, and already shown in similar protocols (Altimus et al., 2008; Tsai et al., 2009; Muindi et al., 2013). Nevertheless, HDC−/−mice exhibited a reduced amount of wakefulness (W) during the biological night but a similar amount during the biological day. Slow wave sleep (SWS) significantly increased during the biological day, the biological night, and over 24h while PS decreased slightly during the biological day and over 24h in HDC−/− mice as compared to their HDC+/+ counterparts. Moreover, sleep-wake amounts during the biological day and night and over 24h in the two mouse strains under the LD1:1 light condition was comparable to that under either the natural light-dark or constant darkness condition, indicating that this ultradian light condition did not significantly disturb the circadian rhythmicity nor its homeostasis in both genotypes (**Fig. 1C and Fig. S2**).

**Figure 1.**
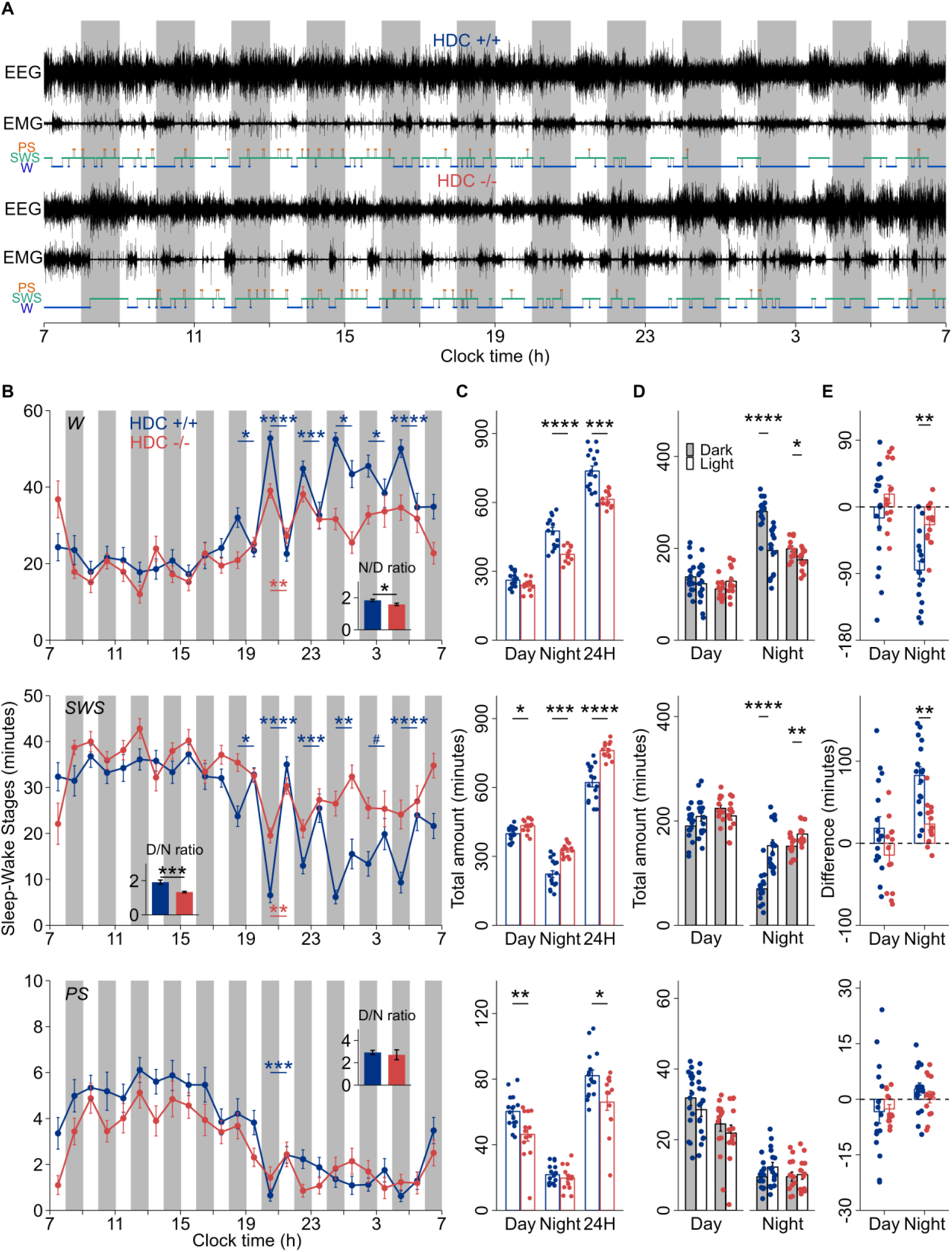
The Sleep-Wake stages under a LD1:1 condition in HDC +/+ (16 tests, blue) and HDC −/− (13 tests, red) littermates. **(A)** Typical examples of polysomnographic recordings and their corresponding hypnograms under the LD1:1 light condition in both genotypic mice (top HDC+/+, bottom HDC−/−). Light schedules are depicted with vertical grey (dark pulse) and white bars (light pulse). **(B)** Time course of mean hourly amounts of W, SWS and PS. Inserted histograms correspond to the biological Night/Day (N/D) or the biological Day/Night (D/N) ratio of the amounts of vigilance states. HDC+/+ induced SWS at the cost of W in response to each light pulse. Light-induced effects on W, SWS just occurred at the beginning of the biological night. **(C)** Total amounts of W, SWS, PS during the biological day, night and over 24h. **(D)** HDC+/+ and HDC−/− mice have more SWS during the six 1-h light pulses than that the six 1-h dark pulses during the biological night (p all<0.05), but not during the biological day. **(E)** Amplitudes of W, SWS, PS changes from dark pulses to light pulses show that HDC+/+ induced more SWS and less W compared to that in HDC−/− (SWS: 83 vs 23 min, p<0.01, W: 85 vs 24 min, p<0.01). Asterisks represent significant differences for post hoc comparisons (*, p <0.05, **, p<0.01, ***, p <0.001, ****, p<0.0001, #: p=0.058). All values represent mean±s.e.m.

In HDC+/+ mice, light pulses significantly increased the amounts of SWS at the expense of W during the biological night, from 70 ± 7 min to 153 ± 11 min (n=8, p<0.0001), but not during the biological day. In HDC−/−, light also enhanced SWS only during the biological night (n=7, p=0.0074), albeit with a dampened response (152 ± 5 min during the dark pulses vs 175 ± 5 min during the light pulses). This suggests that histamine mediates most (~72 %, i.e., 83 min in HDC+/+ vs 23 min in HDC−/−) of the effect of light on sleep-induction observed in HDC+/+ mice. Light did not increase PS duration in both mouse groups (**Fig. 1D-E**).

To further examine whether the EEG quantitative effects of light on sleep and wake were also associated with EEG qualitative changes in specific frequency bands, we analyzed the power spectral density of each behavioral stage during the biological night. In HDC+/+ mice, light elicited a significant increase in delta activity (0.5-3Hz) during SWS at the expense of theta activity (3-9 Hz), indicative of an enhanced sleep pressure and/or quality (n=8, delta: p=0.0017; theta: p=0.0044). A small but significant decrease in spindle rage (9-15 Hz) can be identified during W indicating enhanced alertness, contrast with the deeper SWS. These effects were all absent in HDC−/−mice. Indeed, the identical light pulse did not impact any band across either sleep-wake stages (n=7, p all >0.05). All qualitative aspects during PS were unchanged in both genotypes (**Fig.2**).

**Figure 2.**
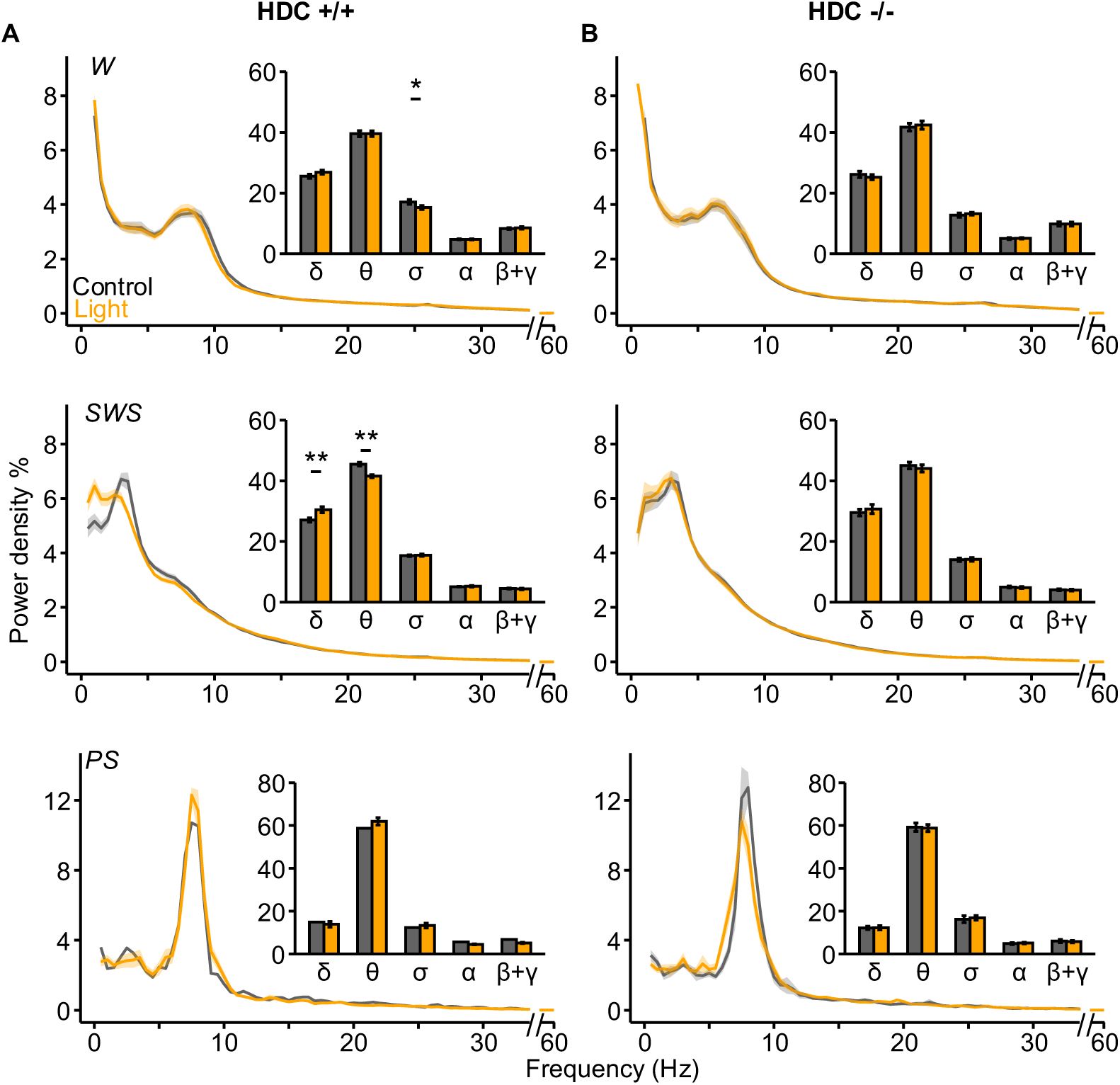
Mean spectral distribution of Cortical EEG power density during W, SWS and PS in HDC +/+ (A) and HDC −/− (B) littermates. Data were analyzed by pooling successive 10 s epochs between 21:00 - 22:00 under the LD1:1 condition (light pulse, orange) and under the constant darkness condition (dark pulse, black) as controls (see Fig. 1B for protocol). To get a better visualization of the low frequency spectral power part, the x axis was broken between 35-55 Hz (no significant difference around this frequency). The inserted bars refer to the EEG spectral power band (δ: 0.5-3 Hz, θ:3-9 Hz, spindle: 9-15 Hz, α:15-20 Hz, β:20-30 Hz, γ: 30-60 Hz) during the light pulse and controls. The peak of spectral frequency during SWS shifts to a lower frequency during light exposure in HDC+/+ mice, but not in the HDC−/− ones. Note that light enhanced delta power and reduced that of theta during SWS while decreasing spindle activity only during W in HDC+/+ mice, but not in HDC−/− mice. Asterisks represent significant differences for the post hoc comparisons (*, p <0.05, **, p<0.01, ***, p <0.001, ****, p<0.0001). All values represent mean ± s.e.m.

### The sleep-promoting effect of light modulated by histamine is time-dependent

To explore whether the effects of light on sleep mediated by HA during the active phase were circadian-dependent, we compared sleep during each dark pulse and subsequent light pulse thorough the biological night in both genotypes. We found that each light pulse significantly enhanced SWS and decreased W in HDC+/+ mice. This sleep-promoting and wake-inhibitory effect of light was especially strong at the first and last 1-h L/D cycle, corresponding to the two peaks of W seen during darkness, characteristic of C57BL/6J mice. In HDC−/− mice, the SWS-enhancing effect of light was observed only during the second 1-h light pulse (21:00, **Fig. 1B**), and this response was smaller versus in HDC+/+ mice, i.e., 11 versus 28 min). Light exerted a PS-enhancing effect just at the beginning of the biological night in association of the high SWS enhancement in HDC+/+ mice, but not in HDC−/− mice (**Fig. 1B**). These findings illustrate a time-dependent feature of the of HA on the SWS-enhancing effect of light, particularly during the beginning and the end of the night.

### The dynamics of sleep and EEG during the light pulse under the LD1:1 condition

To examine how the dynamics of sleep and wakefulness were impacted during light exposure, we looked at the time course of EEG activity every 10 min during the 1-h light pulse occurring at 21:00 (**Fig. 1B**), when the effects of light on sleep were robust, and at a time when both homeostatic and circadian drives for sleep are high in both genotypes. The values of either behavioral stage reached during the dark phase at the equivalent circadian time under DD condition (the day before) served as control. We found a rapid and long-lasting SWS-promoting and W-inhibiting response as early as the first 10 min of light exposure in HDC+/+ mice (W: light: p<.0001, time: p<.0001, light × time: p<.0001; SWS: light: p<.0001, time: p<.0001, light × time: p<.0001), and this response was sustained over the entire duration of the 1-h light pulse (**Fig. 3A**). HDC−/− mice, on the other hand, just showed a delayed and transient induction of sleep after 30 min of light exposure (**Fig. 3B**). We also found a significant enhancement of PS during the second half of the light exposure in HDC+/+ (n=8, PS: light: p<.0001, time: p<.0001, light × time: p<.0001), but not in HDC−/− mice (**Fig. 3A-B**).

**Figure 3.**
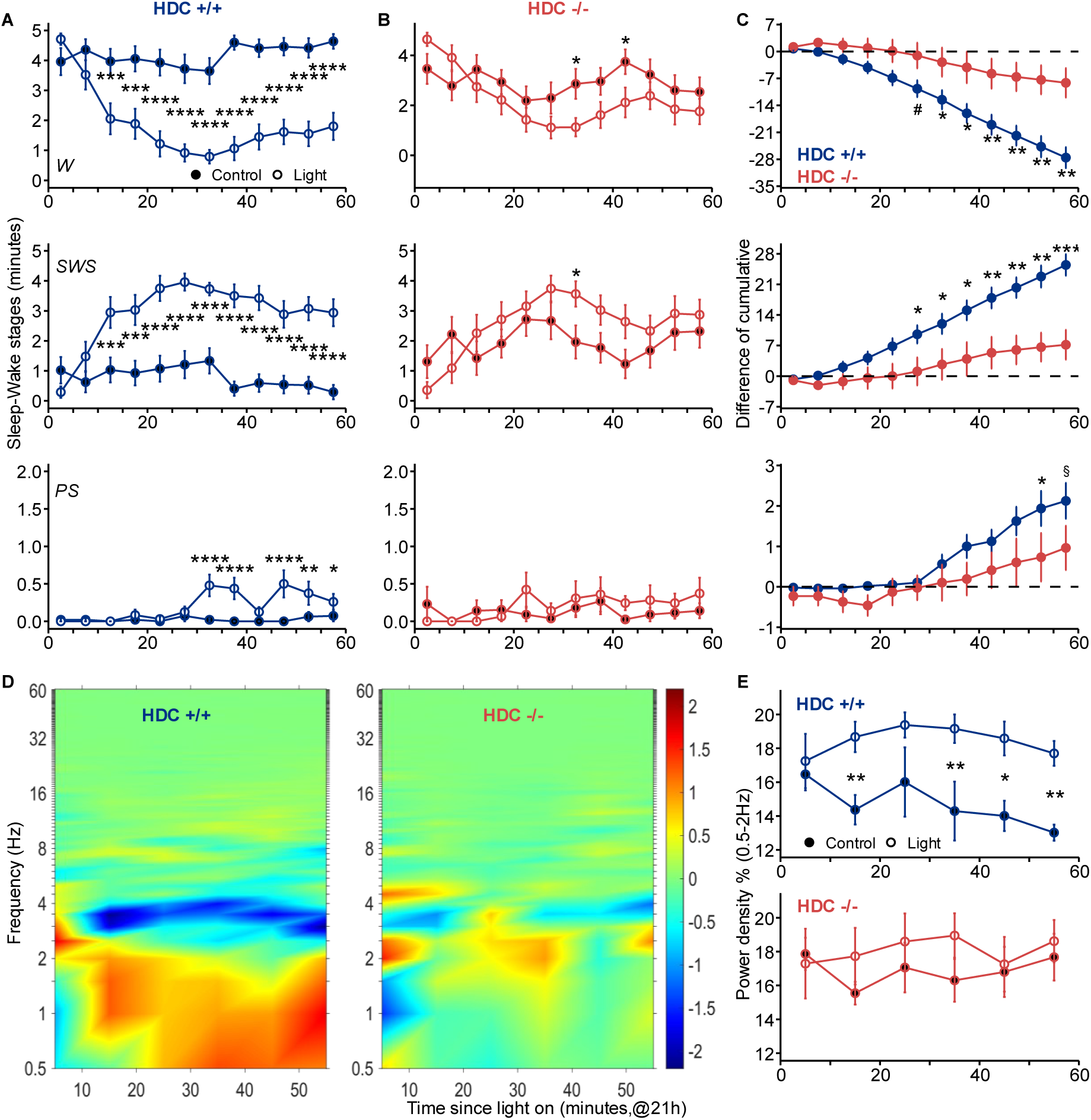
The dynamic of Sleep-Wake stages and the SWS EEG in HDC +/+ (A, 16 tests) and HDC −/− (B, 13 tests) littermates. **(A)** Changes in total amounts of W, SWS, PS between 21:00 - 22:00 (light pulse, unfilled point) under the LD1:1 condition and same clock time under DD condition (dark condition, filled point) in HDC +/+ and **(B)** HDC−/−mice. Data were binned into 5-min intervals. Blue and Red lines represent HDC +/+ and HDC −/− mice, separately. There is a substantial, rapid and sustained sleep-promoting and wake-inhibiting effect induced by light in HDC+/+ mice, while these effects are smaller in HDC−/−, delayed and transient. **(C)** Time course of the accumulated differences during light exposure (deviation from the dark control). HDC−/− accumulated 20 min less SWS during the 1h light pulse compared to HDC+/+ mice. **(D)** Heatmap of EEG changes during SWS during light the 1-h light exposure (deviation from controls); time resolution is 10 min. Warmer colors indicate increases in relative power, while cooler colors represent decreases. **(E)** as in **(D)** summarized for the low-frequency band (0.5 - 2 Hz) shows a reduced and delayed effect in HDC−/− mice. Asterisks represent significant difference for post hoc comparisons (*, p <0.05, **, p<0.01, ***, p <0.001, ****, p<0.0001, #: p=0.054, §: p=0.054). All values represent mean ± s.e.m.

To further investigate the dynamics of sleep induction by light over the 1-h light pulse, we estimated the time course of the cumulative amounts of each sleep-wake stage (deviating from controls) with a time resolution of 5 min in both genotypes (**Fig. 3C**). The accumulation of W or SWS was linear over time during light exposure in HDC+/+ mice, whereas HDC−/− mice showed a much lower accumulation of both states. Overall, HDC−/− mice exhibited a loss of 19 min of SWS during the 1-hour light pulse compared to the HDC+/+ mice, i.e., 7 min versus 26 min. (p=0.0006) (**Fig. 3C**). Altogether, these results suggest that the dynamics of the sleep-inducing light effects depend largely on the HA transmission.

Furthermore, we assessed the dynamics of the qualitative aspects of the light-mediated SWS and found in HDC+/+ mice, that light increased the lower delta frequencies (0.5 - 2 Hz) during SWS, indicative of deeper and more restful sleep, after only 10 min of light exposure, and lasting over the entire light pulse (light: p<.0001, time: p=0.2013, light × time: p=0.1589). In sharp contrast, SWS cortical EEG throughout the light pulse was not affected in HDC−/−mice (n=7, light: p=0.1292, time: p=0.8466, light × time: p=0.7893) (**Fig. 3D-E**) indicating clearly that the SWS quality-enhancing effect of light requires HA transmission.

### Impact of an acute silencing of HA neurons on the sleep-wake effects mediated by light

In order to assess the consequence of an acute silencing of TMn HA neurons on the sleep-wake effects of light, we designed hM4Di-HDC-cre mice by crossing hM4Di and HDC-Cre mice. In this model (**Fig.4A**), HA neurons expressing Cre-recombination are supposed to be acutely silenced by the DREADD ligand DCZ via a modified human M4 muscarinic receptor (hM4Di), thus allowing quantify how an acute single 1h-dark pulse either during the day or 1h-light pulse during the night impact sleep (**Fig. 4**).

**Figure 4.**
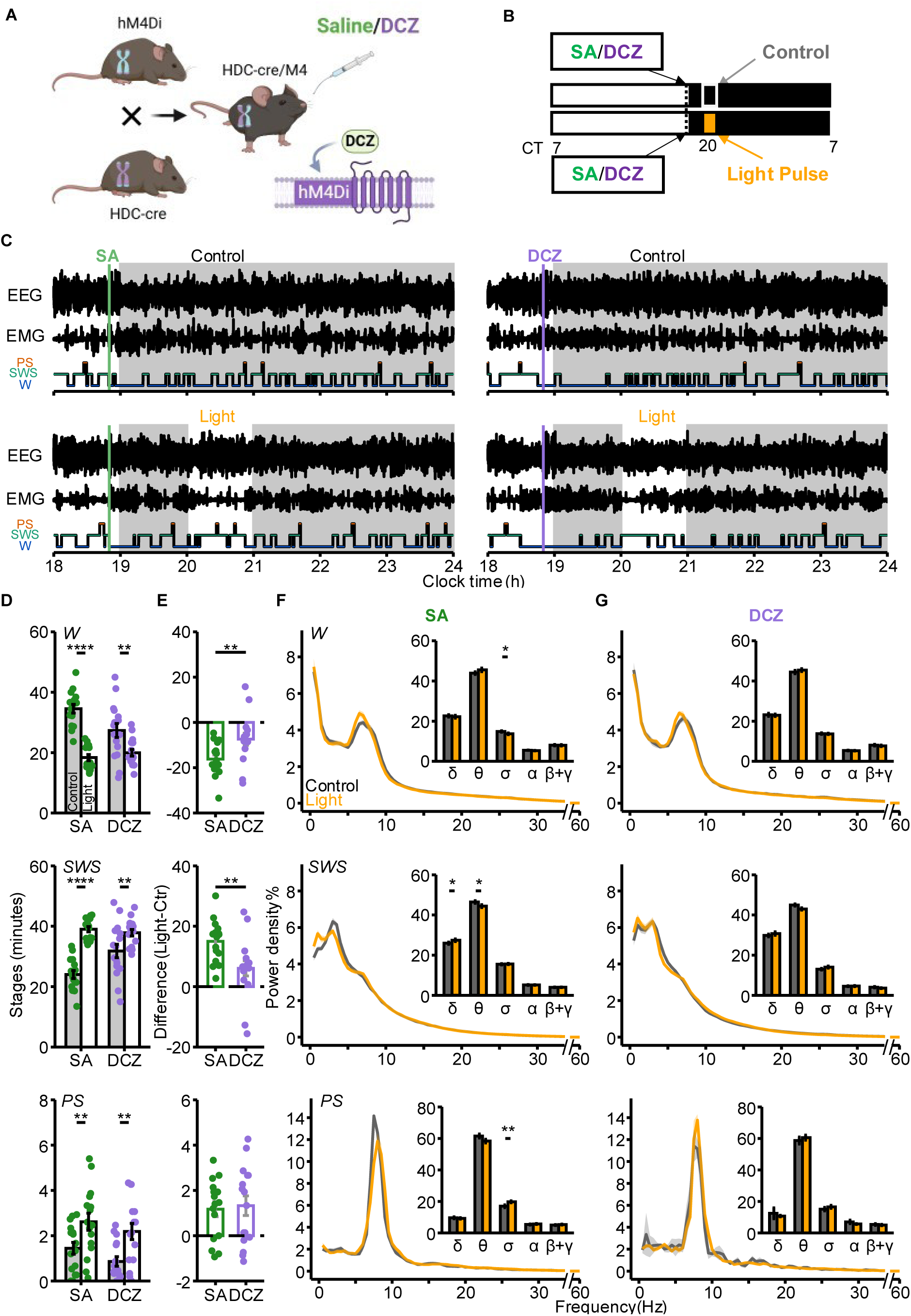
The acute effect of 1-h light pulse during the biological night on the sleep-wake states and the SWS EEG in M4-HDC-cre mice (16 tests). **(A)** The schematic of mouse strains breeding and oral administration of drug handling. **(B)** The diagram of 1-h light pulse exposure and the time of oral administration. Light pulses (orange blocks) were conducted between 20:00 - 21:00, and Saline (SA) or Deschloroclozapine (DCZ) were administered at 18:50 (dash line), CT: clock time. **(C)** Typical examples of polysomnographic recordings and their corresponding hypnograms under the SA/DCZ conditions. Light schedules are depicted with vertical grey bars (dark phase) and white bars (light phase). The vertical-colored line indicates the administration time. Four conditions in the same mouse: Saline group (21 min vs 41 min) and DCZ group (28 min vs 35 min) for control vs light conditions, respectively. **(D)** The acute effect of the 1h-light pulse on the total amount of sleep in SA (green) and DCZ (purple) group. The grey bar represents the data from the control condition and the white bar indicates the data from the light pulse. **(E)** The difference of total amounts of sleep states between the light pulse and control. Light pulse significantly induced SWS (15 vs 6 min, p=0.0016) and decreased W (16 vs 7 min, p=0.0028) either in SA or DCZ administration group, but with a smaller amplitude after DCZ administration group. **(F)** and **(G)** Mean spectral distribution of Cortical EEG power density during W, SWS and PS during the period of light exposure and controls after administering SA and DCZ. To get a better visualization of low frequency spectral power part, the x axis was broken between 35-55 Hz (no significant difference around this frequency). The inserted illustration refers to the EEG spectral power band (δ: 0.5 - 3 Hz, θ:3 - 9 Hz, spindle: 9 - 15 Hz, α:15 - 20 Hz, β:20 - 30 Hz, γ: 30 - 60 Hz) during light and control. Black represents the darkness controls and orange represents the light. Note that the peak of spectral frequency in SWS shifts to lower frequency during light pulse compared to controls after administrating SA but not after DCZ administration. Light decreased spindle activity of W and theta activity of SWS, decreased theta power of SWS after SA administration, but there is no significant difference between light and controls after DCZ administration. Asterisks represent significant difference for post hoc comparisons (*, p <0.05, **, p<0.01, ***, p <0.001, ****, p<0.0001). All values represent mean ± s.e.m.

#### Effects of a 1-h dark pulse during the biological day

In a LD12:12 condition (**Fig. S1A**), light was switched off from 12:00 to 13:00 with oral administration of saline or DCZ one hour before. The dark pulse did not significantly impact sleep and W in the Saline and DCZ group. There was no significant impact of the dark pulse on any band of spectral power in two groups (**Fig. S1B-E**).

#### Effects of a 1-h light pulse during the biological night

When light was switched on at the beginning of the night (from 20:00 - 21:00) with oral administration of saline or DCZ one hour before, we found that, in sharp contrast to the day test, light remarkably induced SWS and decreased W in the saline group (**Fig. 4D**, p all <0.0001). In the DCZ group, SWS increased and W decreased during the light pulse (**Fig. 4D**, control vs light: SWS, 32 vs 38 min, p=0.0062; W, 27 vs 20 min, p=0.0025) but with smaller magnitude regarding either SWS or W compared to the saline group (**Fig. 4E**, SWS induced by light: 6 vs 15 min, p=0.0016; W inhibited by light: 7 vs 16 min, p=0.0028).We also found a significant increase in PS duration during the light pulse in both saline and DCZ groups, with a similar amplitude (n=8, p=0.811) (**Fig. 4D, E**). Regarding the qualitative aspects, there was a light-induced increase in the spindle band (9 - 15 Hz) during W, in the saline group (n=8, p=0.0466, **Fig. 4F**), but not in the DCZ group (n=8, p=0.8780). Notably, light considerably increased delta activity (0.5 - 3 Hz) at the expense of theta activity (3 - 9 Hz) during SWS in the saline group (n=8, delta: p=0.0481; theta: p=0.0479). These responses to light were absent after DCZ administration (n=8, p all >0.05, **Fig. 4G**). An enhancement of spindle activity (9 - 15 Hz) during PS was observed during the light pulse in the saline group (n=8, p=0.0053), but there was no difference in any spectral bands between light pulse and controls during PS after administering DCZ (n=8, p=0.4342) (**Fig. 4G**). These results in hM4Di-HDC-cre mice reveal the major role of HA transmission in the acute photic modulation of the sleep EEG.

### Acute light effects on the dynamics of sleep and EEG

To examine how the dynamics of sleep and W were impacted during light exposure upon silencing HA transmission, we looked at the time course of EEG changes every 10 min during the 1-h light pulse (**Fig. 5**). We found that light induced a significant decrease in W and an increase in SWS after 10 min of light exposure in the saline group, consistent with what we found under the LD1:1 condition (n=8, W: light: p<.0001, time: p=0.5216, light × time: p=0.0083; SWS: light: p<.0001, time: p=0.2368, light × time: p=0.0133). However, in the DCZ group, the response to light pulse was delayed and transient compared to baseline (n=8, W: light: p=0.0008, time: p=0.0236, light × time: p=0.0002; SWS: light: p=0.0042, time: p=0.0441, light × time: p=0.0031). The effect of light on PS was comparable in the two groups, with both showing a transient increase in PS over the acute light pulse (Saline: n=8, light: p=0.0115, time: p=0.2899, light × time: p=0.0359; DCZ: light: p=0.0041, time: p=0.7988, light × time: p=0.1738) (**Fig. 5A-B**). Analyzing the cumulative amounts of each sleep stage over the 1-h light exposure shows that mice administered DCZ accumulated about 9 min less SWS compared to the saline group (**Fig. 5C**). This confirms that light induces sleep with a quick, persistent and substantial response and requires HA transmission.

**Figure 5.**
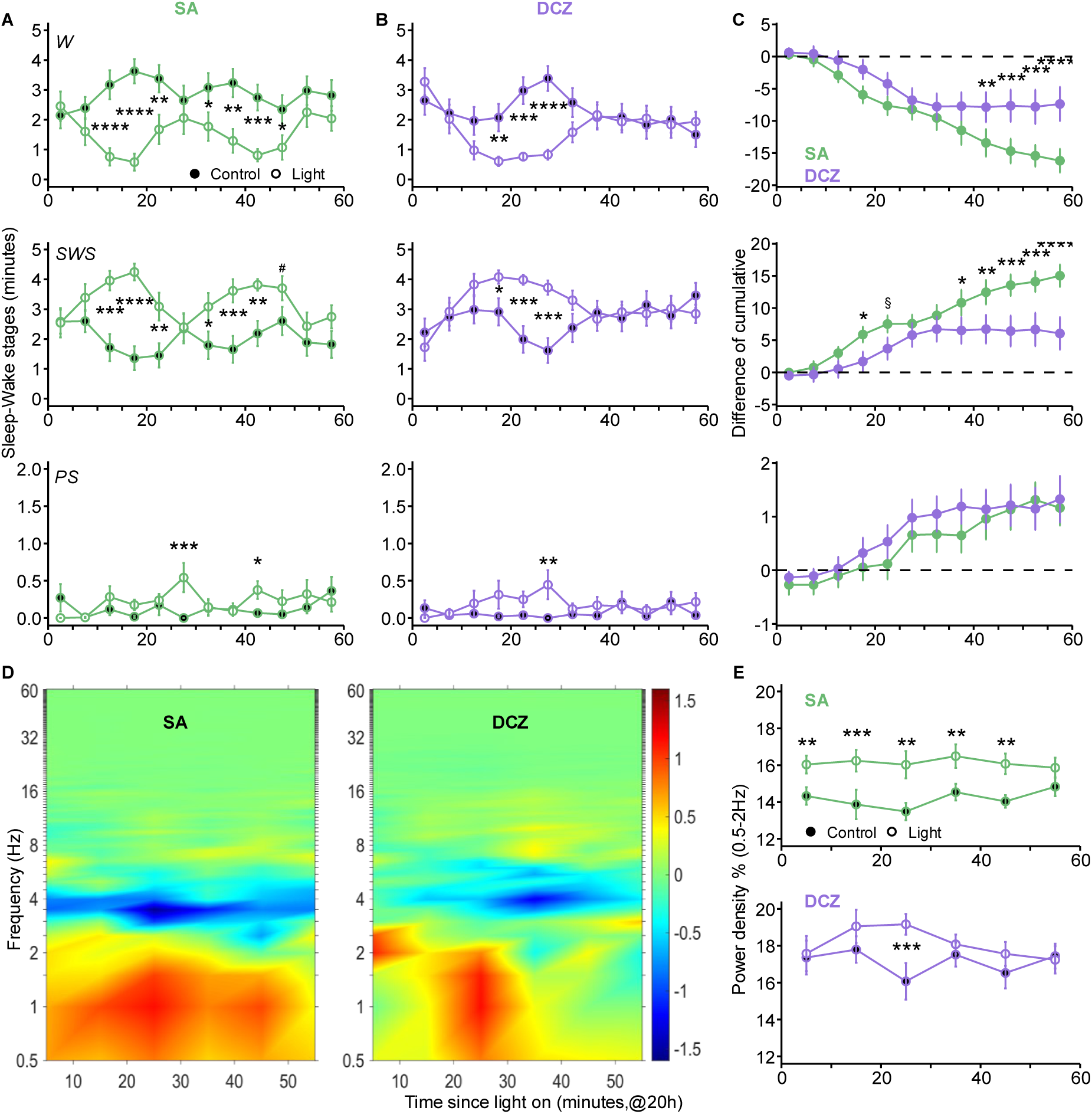
The dynamics of Sleep-Wake stages and the SWS EEG during the 1-h light pulse period in hM4Di-HDC-cre mice (16 tests). **(A)** Changes in total amounts of W, SWS, PS between 20:00-21:00 (light pulse, unfilled point) and same clock time under a standard LD condition (dark condition, filled point) after SA and DCZ administration **(B).** Data were binned into 5-min intervals. Green and purple lines indicate SA and DCZ groups, separately. There is a substantial, rapid and sustained sleep-promoting and wake-inhibiting effect induced by light in SA group, while these effects are smaller in DCZ administration group. **(C)** Time course of the accumulated differences during light exposure (deviation from the dark control). During the 1-h light exposure, DCZ group accumulated 10 min less SWS compared to SA administered group. (**D**) Heatmap of EEG changes during SWS during light the 1-h light exposure (deviation from controls); time resolution is 10 min. Warmer colors indicate increases in relative power, while cooler colors represent decreases. **(E)** as in **(D)** summarized for the low-frequency (0.5 - 2 Hz) bands demonstrate a reduced and delayed effect in DCZ administered group. Asterisks represent significant difference for post hoc comparisons (*, p <0.05, **, p<0.01, ***, p <0.001, ****, p<0.0001, #: p=0.054, §: p=0.057). All values represent mean ± s.e.m.

Looking into whether light also impacted brain activity during SWS and the role of HA therein, we found that the light-induced SWS was characterized by an immediate and long-lasting increase in the EEG low delta frequencies (0.5 - 2 Hz) after saline administration (light: p<.0001, time: p=0.8559, light × time: p=0.7467). In contrast, in the DCZ group, only a temporary light-induction of delta activity was noted (light: p=0.0105, time: p=0. 3883, light × time: p=0.1188). (**Fig. 5D-E**), suggesting that the dynamics of sleep depth are altered following silencing HA neurons.

### Dual Immunohistochemistry for HDC and HA-Tag

At the end of sleep-wake experimentation and to evaluate the recombination rate between HA neurons and the hM4Di receptor, hM4Di-HDC-cremice were perfused for dual immunohistochemistry of histidine decarboxylase (HDC), marker of histamine neurons, and HA-Tag, marker of the hM4Di receptor. Confocal microscopic observation indicated that a great number of HDC-immunoreactive (-ir) cells are HA-Tag positive (**Fig. 6**), indicating recombination of major HA neurons with hM4Di receptor following crossing between HDC-cre and hM4Di mice. Cell counts further revealed a strong expression of HA-Tag int 69% of HDC-ir neurons. This recombination rate appeared unexpectedly relatively moderate. On the one hand, our cell count was performed with low microscopic magnification, only strongly visible labeling was considered while weakly labeled but functionally recombinant cells might be escaped. On the other hand, low range concentrations of antibodies were used to ensure the specificity of the labeling but to some extent might have compromised the immunohistological sensitivity. Nevertheless, in view of the clear phenotypes identified in this study using light as stimulus and in other behavioral tests (Wu et al., n.d.) that are highly consistent with the findings obtained from knockout models, we can affirm here that this recombination rate was underestimated but functionally suitable for validation of our experimental model.

**Figure 6.**
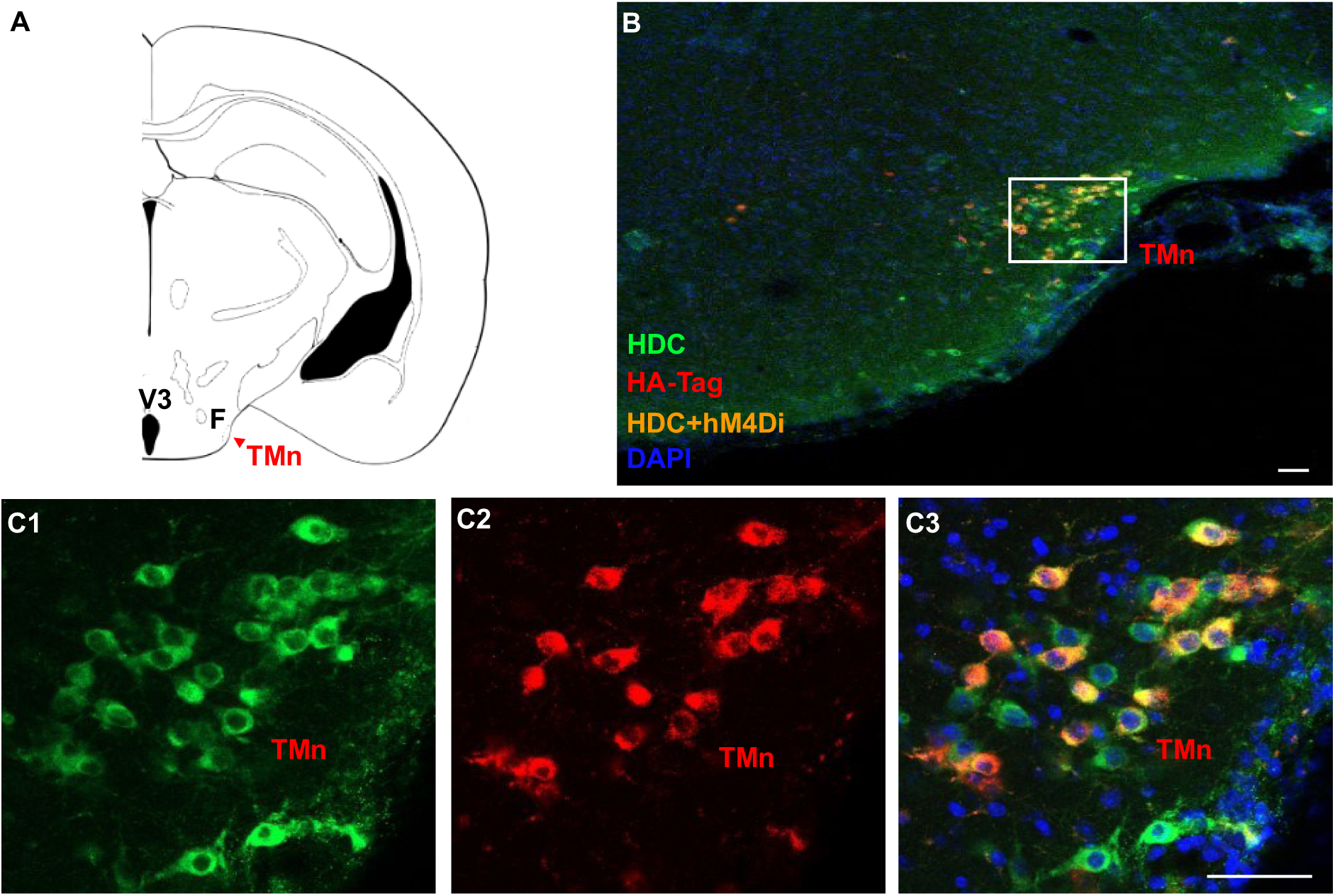
Expression of hM4Di-receptor labeling in HDC immunoreactive tuberomammillary histaminergic neurons (TMn) in hM4Di-HDC-cre mice. **(A)** Schematic of a coronal section of the brainstem showing the location of the TMn. **(B)** A confocal photomicrograph of lower magnification showing both histidine decarboxylase (HDC (green) and HA-Tag (red) single and double (orange) fluorescent labeling in the TMn and adjacent posterior hypothalamus. **(C1-C3)** photomicrographs of higher magnification of the boxy insert in B showing the number and morphology of HDC-immunoreactive (-ir, green), HA-Tag positive (red) and dual-labeled (orange) cells in the TMn. Note that a large number of HDC-ir cells are HA-Tag positive, indicating recombination of major histaminergic neurons with hM4Di receptor following crossing between HDC-cre and hM4Di mice. Other abbreviations: 3V, third ventricle, F, fornix. White scale bar =100 μm.

## Discussion

The effects of light on sleep can be indirect—via photo-entrainment of the circadian system— or direct, through non-circadian pathways. Our study reveals that HA neurons, traditionally associated with wake promotion, mediate the direct sleep-inducing effect of light. Specifically, we found that, only during the biological night, light not only increased the amount of SWS, but also cortical delta activity —an EEG marker of sleep depth. These effects were abolished or markedly attenuated in mice with long-term loss of HA transmission or following acute silencing of TMn HA neurons.

### The effects of light on sleep and histamine neurotransmission

The acute effects of light on sleep in rodents have been well documented (Borbély et al., 1975), with ipRGCs in the retina identified as a pivotal factor (Altimus et al., 2008; Tsai et al., 2009). The neural pathways responsible for this effect have been very partly identified. A study reported that the lesion of the superior colliculus (SC) of the midbrain, a region that modulates visual processing, orientation, and eye movement, resulted in the elimination of the acute changes in sleep pattern in response to light observed in albino rats (Miller et al., 1998). Another study showed that the intergeniculate leaflet (IGL) of the thalamus, which is responsible for visual motor and circadian time-keeping systems, was also involved in light-induced sleep in mice (Shi et al., 2020). More recently, the acute effects of light on sleep were clearly shown to be mediated by the pathway from the retinal ipRGCs to the preoptic area (POA) (Zhang et al., 2021). In that study, the POA neurons were additionally identified as forming inhibitory monosynaptic connections with wake-promoting regions, including the TMn, the LH, the ventral tegmental area (VTA), and the dorsal raphe nucleus (DRN) (Zhang et al., 2021). However, none of these downstream structures were investigated during sleep induction by light and therefore their functional involvement and relative contribution remains unknown. Our study was designed to bridge this gap by assessing the sleep induced effect of light upon short- or long-term deletion of HA transmission.

Phenomenologically speaking, light induced a greater enhancement of SWS in HDC+/+ than in HDC −/− mice. Was this due to the fact that HDC −/− mice were more asleep during their baseline condition (Anaclet et al., 2009) and so presented less opportunity for SWS enhancement during light exposure? This is unlikely, because our mice slept with eyes open as previously reported (Yüzgeç et al., 2018). In addition, despite the variable pupil constriction during sleep and W, we calculate that sufficient light still reaches the retina to induce non-visual responses at the levels mice were exposed to in our study (Zhang et al., 2017). This leads us to exclude that the reduced sleep-inducing effects observed in HDC −/− mice results from a decreased light input due to pupil size or eye closed.

The 72% reduction in light-induction of SWS in HDC−/− mice speaks for the primary role of HA in this effect. Thus, HA neurotransmission may account for most of the effects (~70%) of light on sleep in control animals, while other pathways may account for only a minor part (~30%). The other structures innervated downstream of the ipRGC-POA pathway, such as the LH, the VTA, and the DRN as well as, possibly, the locus coeruleus (LC), are potential contributors to this remaining, non-histaminergic effect observed in HDC −/− mice.

It has been shown that chronic KO models embed up- or down-regulation in different neural systems to offset the loss of a specific pathway. Therefore, it cannot be excluded that the effects on light-induced sleep observed in the HDC −/− model result from such compensatory mechanisms. To address this, we included a study using a Cre-dependent model coupled with the DREADD system, allowing for acute and reversible silencing. We employed the novel DREADD ligand DCZ at a low dose (0.5 mg/kg) to ensure pharmacogenetic specificity and found that acute silencing of TMn HA neurons significantly reduced the SWS-promoting effect of light and abolished the associated increase in delta power—effects that were rapid, sustained, and circadian-specific, occurring only during the biological night, highly similar to those seen with HDC −/− mice. The consistency of the results found in acute and chronic models validates the findings in both approaches and suggests that certain key phenotypes are robust and not eliminated by the often-powerful brain compensatory plasticity.

The SWS enhancement observed in our study cannot be readily explained by a light-induced inhibition of TMn HA neurons via ipRGC/POA inputs, as proposed by Zhang et al. (2021). Indeed, various experiments impairing or inhibiting TMn or HA transmission in WT models consistently enhance SWS - fundamental evidence allowing the classification of HA as a “waking amine” (J. C. Schwartz et al., 1991; Haas et al., 2008; Lin et al., 2011; J.-C. Schwartz et al., 2011; Arrigoni & Fuller, 2022). However; our results showed the opposite effect under light exposure in mice, in which HA is not merely inhibited but completely abolished or silenced. These surprising findings thus challenge the traditional view of HA as W-promoting transmitter and highlight the requirement of a normal HA functioning for light-induced SWS rather than its absence or inhibition. One might also speculate that in KO mice, the absence of HA in the TMn could be replaced by other transmitters such as GABA, HA’s co-transmitter. This could potentially transform TMn neurons into inhibitory sleep-promoting cells, whose inhibition by ipRGC/POA inputs would paradoxically result in sleep loss. However, this remains entirely hypothetical in the absence of direct experimental evidence.

Alternative pathways should be considered. Notably, HA receptors—H1, H2, and H3—are expressed in the retina (Sawai et al., 1988; Gastinger et al., 2006; Greferath et al., 2009; Vila et al., 2012), where they may modulate retinal sensitivity to light or gate retinal inputs to brain regions controlling circadian rhythms and behavioral states. This could help explain, at least in part, the decreased light-induced sleep response observed in HA-deficient mice.

HA has primarily been studied in the context of visual processing (Gastinger et al., 2004; Yu et al., 2009; Akimov et al., 2010; Tripodi & Asari, 2024; Warwick et al., 2024), where it influences retinal ganglion cell (RGC) activity in a subtype-specific manner. Whether HA exerts a similar modulatory role in non-visual retinal functions remains to be determined. In particular, it is unknown whether retinal HA can regulate the activity of ipRGCs, especially the Brn3b-positive subtype known to mediate light-induced sleep (Rupp et al., 2019; Tran et al., 2019).

In the face of the major challenge of understanding how a wake-promoting system can be recruited for light sleep induction, the priority should be identifying the underlying neural circuit responsible for these surprising findings. In particular, investigate whether retinal inputs reach brain areas responsible for cortical activation and W such as the TMn, and conversely, whether TMn HA neurons send direct projections to the retina, in addition to the local expression of HA receptors. At this point, there is no evidence of direct projections of ipRGCs to W-promoting structures (Hattar et al., 2006; Morin, 2015), but more sensitive approaches should be used before ruling them out.

### Time- and circadian-dependent regulation of light induced sleep and histamine transmission

Our results confirm that the sleep-inducing effects of light are gated by the circadian system, as they were observed during the biological night (habitual period of darkness) and not the biological day (habitual period of light exposure), both in control and HA-deficient mice. These characteristics are linked with the circadian rhythmicity of HA transmission. In nocturnal rodents, HA release is indeed high during darkness and low during lightness with a peak just before and after lights-off (Mochizuki et al., 1992). Pharmacological inhibition of HDC enzyme enhances sleep only during darkness (Parmentier et al., 2002; Gondard et al., 2013). HA-deficient models or silencing TMn HA neurons display sleepiness only around lights-off when the brain requires HA (Parmentier et al., 2016; Wu et al., n.d.), while the sleep-promoting POA/VLPO is maintained at its lowest (Gvilia et al., 2006), by the tonic inhibition of W promoting transmitters (Gallopin et al., 2000), including HA (Liu et al., 2010), through the recently discovered ipRGC-POA pathway and projections to the wake promoting structures (Zhang et al., 2021). Conversely, the absence of a light effect during daytime could result from the minimal activity of the TMn at that circadian time, and therefore the incapacity of light to further suppress the wake-promoting HA system. This in no way claims that HA is the only neurotransmitter involved in the sleep-induction by light. On the contrary, we and others (Zhang et al., 2021) suggest that other pathways are involved, in a dynamic, non-binary fashion (biological day versus biological night). Indeed, Orexin knockout mice show similar phenotypes although in a slightly less extent than that of HDC −/− mice regarding light-induced SWS (Tang et al., n.d.).

### The effects of light are dynamic, rapid, sustained

As previously shown, the effects of light on sleep-wake regulation are rapid, with locomotor suppression and subsequent sleep onset after only two minutes of light exposure, which persist for approximately 20 minutes in rodents (Muindi et al., 2015). In humans, the non-visual effects of light exposure activate EEG, heart, pupillary and thermoregulatory responses within one to five minutes (Prayag, Jost, et al., 2019). Our study shows that light can induce sleep within 10 minutes, and in a sustained manner though the entire light pulses, in the control animals but not the HA-deficient animals where the response is delayed and reduced. The process of inducing sleep through light exposure depends on the interaction between neurotransmitter systems that promote sleep and those that hinder Ws, as well as modulation by the circadian timing system. Overall, our results underscore the importance of HA in mediating a quick and long-lasting response to light.

## Conclusions

In conclusion, our findings indicate that HA neurotransmission plays a primary role in mediating the acute effects of light on the rapid and sustained induction of SWS, and that the acute sleep-inducing effects of light require HA transmission in mice.

We propose that the circadian-gated effect of light relies mainly on the circadian regulation of HA transmission, and that the absence of light-induced sleep during the day, is not, as classically discussed, due to the impossibility to produce more sleep during the day (ceiling effect), but to the incapacity of light to further inhibit wakefulness (floor effect).

## Methods

### Animals

This study used adult C57BL/6J mice with an age range of 13-26 weeks including 8 drug-naive hM4Di-HDC-cre (served as HDC+/+ or wild type or control), 7 histidine decarboxylase gene knock-out (HDC−/− or KO) (Study 1) and another 8 hM4Di-HDC-cre mice (Study 2). HDC−/− mice were generated according to previously described procedures (Ohtsu et al., 2001). hM4Di-HDCcre mice were generated in our laboratory by crossing hM4Di and HDC-Cre mice through homologous recombination in embryonic stem cells and replacing the HDC gene with Cre recombinase (Fujita et al., 2017). Polymerase Chain Reaction (PCR) was performed on genomic DNA from tail biopsies of all mice to confirm their genotypes. All experiments followed EEC Directive (2010/63/EU) and every effort was made in accordance with 3R principles to minimize the number of animals used and any pain and discomfort. The general protocol of chronic sleep-wake recordings in mice was approved by the local and national Ethic Committee of Animal Experimentation.

### Surgery, housing and polysomnographic recordings

Animals were chronically implanted, at the age of 10-12 weeks under deep anesthesia and painless conditions (Ketamine, 1 μL/g; 2% Xylazine, 0.5 μL/g; Lurocaine, 0.3 μL/g; Carprofen, 0.1 μL/g. i.p.), with five cortical electrodes and two muscle electrodes to record the electroencephalogram (EEG) and the electromyogram (EMG). All electrodes were previously soldered to a multi-channel electrical connector, and each was separately insulated with a covering of heat-shrinkable polyolefin/polyester tubing. The cortical electrodes were inserted into the dura through 3 pairs of holes (Ø ¼ 0.3 mm) made on the skull, located, respectively, in the frontal (1.5 mm anterior to the Bregma and 1 mm lateral to the midline), parietal (2 mm posterior to the Bregma and 1.5 - 2 mm lateral to the midline), and cerebellum (Slightly offset at the midline, slightly closer to the lambda). The 2 muscle electrodes were guided down the back of the neck underneath the trapezius muscles.

Mice were then housed individually in transparent barrels (Ø 20 cm, height 30 cm) within a soundproofed recording room with a constant ambient temperature at 24°C (±2°C), humidity at 50% (±10%) and 12h light/dark cycle (1,000 lux/ 0ux, light on at 07:00 and off at 19:00), food and water available ad libitum.

Polysomnographic recordings and data collecting started after a period of three weeks for recovery from surgery and habituation to the experimental ambiance. Cortical EEG and EMG signals were amplified, digitized with a resolution of 256 and 128 Hz, respectively, and computed on a CED 1401 Plus (Cambridge, UK). The data were analyzed using Spike2 and SleepSign (v1503, Kissei-Comtec, Japan) softwares at 10 seconds (study 1) and 4 seconds (study 2) intervals for sleep stages and power spectral density analysis. The sleep-wake states were scored by experienced staff blind to conditions and genotypes, according to the standard criteria of the laboratory for wakefulness (W), slow-wave sleep (SWS), and paradoxical or REM sleep (PS). EEG power spectra were computed for consecutive epochs within the frequency range of 0.5-60 Hz using a fast Fourier transform routine at 0.5 Hz resolution. The following bands were used in this study: delta (δ): 0.5-3 Hz, theta (θ): 3-9 Hz, (spindle, represented as σ): 9-15 Hz, alpha (α): 15-20 Hz, beta (β): 20-30 Hz, gamma (γ): 30-60 Hz, and beta + gamma (β+γ): 20-60 Hz. To standardize the data, all power spectral densities at the different frequency ranges were expressed as a percentage relative to the total power (e.g., power in the δ band/power in the 0.5-60 Hz) of the same epochs.

### Chemical compounds and pharmacological administration

The DREADD ligand Deschloroclozapine (DCZ; E1247-3-2, HelloBio, Meath, Republic of Ireland) was dissolved in 0.05 mL saline (0.9% NaCl) containing 1% methylcellulose (039K0146, Sigma-Aldrich, St. Quentin Fallavier, France). The DCZ solution was gently administered to each mouse by oral route using a probe for mouse oral application (20G, Phymep, Paris, France) at a dose of 0.5 mg/kg (compound weight), at 10:50 and 18:50, respectively, according to study 2 (see **Fig. 4B** and **Fig. S1A**).

### Light schedules and protocols

Light exposures were conducted with white fluorescent light at 1,000 lux, using two panels leaning against the two adjacent white walls in front of the platform on which mice transparent barrels were placed, so that the horizontal light intensity at the eye level of the mice was homogeneous across animals. The light intensities and spectra were continuously recorded with a spectral photometer (BTS2048-VL, Gigahertz Optik GmbH, München, Germany).

Mice were adapted to a normal 12:12 hours light-dark cycle (LD12) one week before the experiment and habituated to the EEG/EMG cable before starting the recordings. The following protocols were repeated twice.

**1)** 8 hM4Di-HDC-cre mice (HDC+/+) and 7 HDC−/− mice respectively followed a consecutive 3-day schedule including a standard day (LD12, light between 07:00 - 19:00, darkness between 19:00 - 07:00), a day in constant darkness (DD, light off during 07:00 - 07:00), and an ultradian day with alternating 1-h light and 1-h dark pulses throughout the 24 h (LD1:1, light on at 07:00, off at 08:00, on at 09:00, etc.). Light was automatically controlled by a programmable timer switch (Novkit) and no entry in the mice room was allowed during the entire 3-day recording protocol.
**2)** A second group of 8 hM4Di-HDC-cre mice were employed in a 4-day cross-design protocol. Mice were exposed to a standard day (LD, light between 07:00 - 19:00, darkness between 19:00 - 07:00), while with an oral application of Saline (SA) or DCZ at 10:50 or at 18:50 following a light or dark pulse (see **Fig. S1A and Fig. 4A-B**). Mice were kept under an additional LD12 condition after each application for a 1-day washout time.

### Immunohistochemistry of HA-Tag and HDC

#### Preparation of brain sections

At the end of sleep-wake experimentation, hM4Di-HDC-cre mice were anesthetized with ketamine and pentobarbital (both 100 mg/kg, i.p.) and perfused via the left ventricle into the ascending aorta with 50 ml saline followed by ice-cold 4% paraformaldehyde in 0.1 M phosphate buffer (PB). Brains were carefully removed, post-fixed in the same fixative overnight and then immersed in 30% sucrose solution in 0.1 M PB at 4 °C for > 48h. They were coronally sectioned (40 μm) on a cryostat (Leica Micro-systems, Heidelberg, Germany) at −20 °C and collected into cryoprotective solution (30% ethylene-glycol; 20% glycerol; 50% 0.01 M PB) and stored at −20 °C until use.

#### Dual-immunofluorescence procedures

Floating sections were rinsed in 0.01 M PBS solution containing 0.1% Triton X-100 (PBST) for 10 min × 3 times, then incubated in blocking solution (10.0% fetal bovine serum in 0.01 M PBST) for 1.5 h. Sections were incubated with a rabbit monoclonal antibody against HA-Tag (C29F4) (1:1,000 in blocking solution, #3724, Cell Signaling Technology, Massachusetts, USA) for 24 h at 4 °C. After rinsing in PBST, sections were incubated with CyTM3 AffiniPureTM donkey anti-rabbit IgG (1:2,000, 711-165-152, Jackson Immuno Res. Lab, PA, USA) for 5h at room temperature. Rinsed sections were then incubated with a rabbit polyclonal antibody against histidine decarboxylase (HDC) (1:1,000 in blocking solution, 16045, Progen Biotechnik GmbH, Heidelberg, Germany) for 48h at 4°C. After rinsing in PBST, sections were incubated with Alexa Fluor488® AffiniPureTM donkey anti-rabbit IgG (1:2,000, 711-545-152, Jackson Immuno Res. Lab, PA, USA) and DAPI (1:2,000, D8417, Sigma-Aldrch, Saint Louis, MO, USA) for 2h at room temperature. Finally, sections were mounted on slides, covered with a coverslip via 70% glycerol in PBS and sealed with nail polish. Images were captured with the confocal fluorescence microscope (Tissue FAXS PLUS, Tissue Gnostics, Vienna, Austria).

### Statistical analysis

Data and statistical analyses were carried out using Matlab (v. R2020b, MathWorks) and R v.4.3.0 (RStudio 2024.04.2 Build 764) (R Core Team, 2024). Comparisons of sleep amounts or quantitative EEG powers between control and experimental groups were performed using a linear mixed model method (package ‘*nlme’*, v 3.1.164) (Pinheiro, J et al., 2023). And all datasets with multiple comparisons were analyzed by a linear mixed model, followed by a Bonferroni post hoc test. Each analysis was computed after a Box-Cox transformation (package ‘car’, v 3.1.2) to normalize the data with the optimal power transformation. P< 0.05 was considered as the statistically significant threshold. All data are presented as mean ± s.e.m.

## Acknowledgements

Y.H and N.T were supported by a PhD Fellowship from the Chinese Scholarship Council (CSC202006180028 and CSC202006180019, respectively). Funding was obtained by JS Lin and C Gronfier (Inserm, ANR Histawake & Labex Cortex).

## Author contributions

C.G. and J-S.L. conceived the project. Y.H. and N.T. conducted the study, computed all the analyses, produced graphic illustrations and drafted the article. S.W. developed the hM4Di-HDCcre mouse model, contributed to the implantation of electrodes for sleep-wake monitoring, and helped with the initial analysis of the EEG data. Y.S. contributed to the immunohistochemical work.

