## Supplemental figures for "The acute sleep-inducing effects of light require histamine neurotransmission in mice"

* Corresponding author

**Corresponding Authors:**

**Supplementary figures**

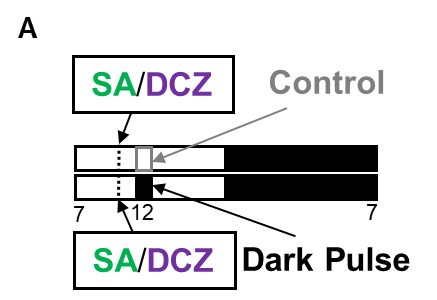

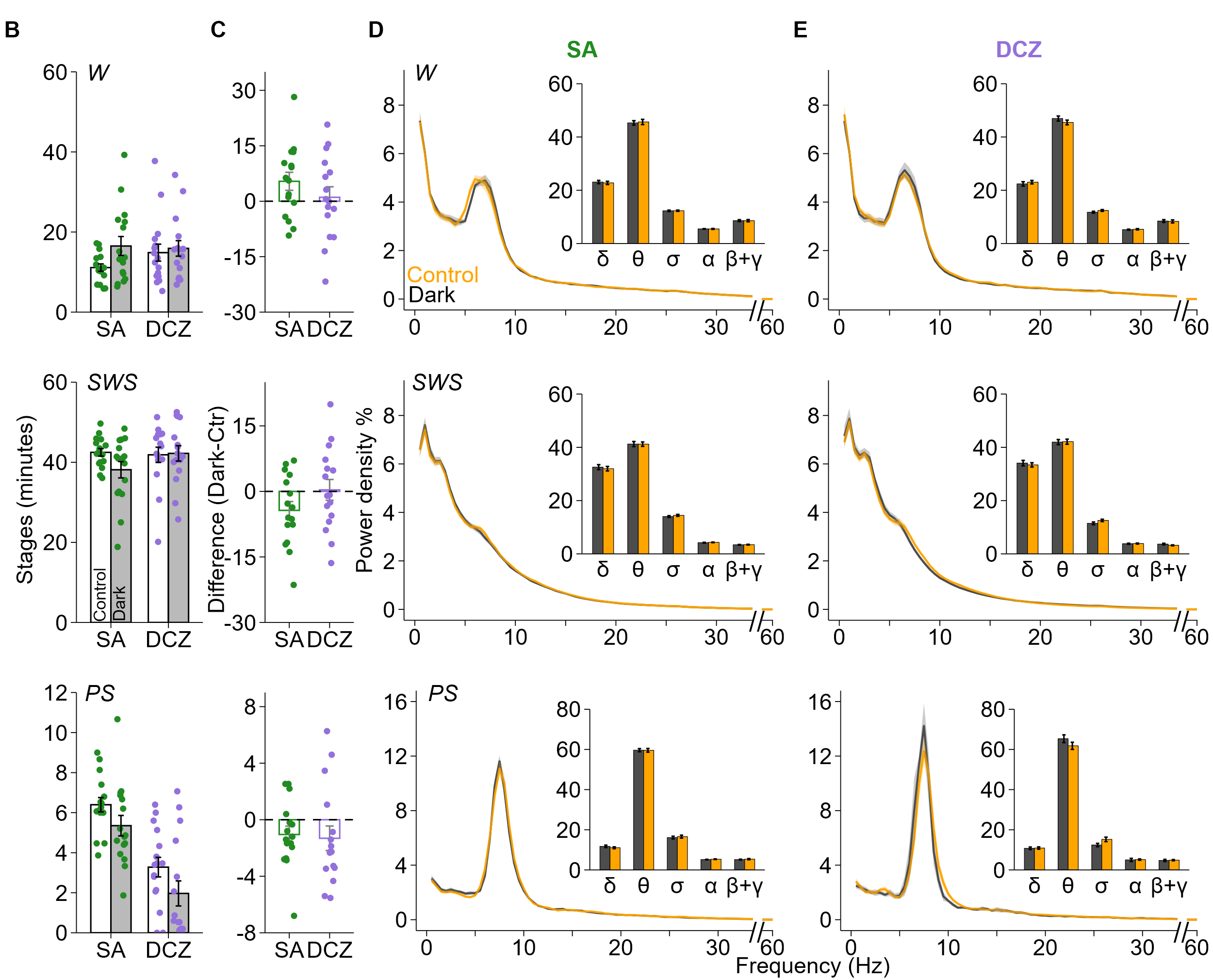

**Figure S1 The acute effect of 1-h light pulse on the sleep-wakefulness distribution and the SWS EEG during the biological day. Left top showed the light protocol and time for administration. (A)** The diagram of 1-h light pulse exposure and the time of oral administration. Light pulses (orange blocks) were conducted between 12:00 -13:00, and Saline (SA) or Deschloroclozapine (DCZ) were administered at 10:50 (dash line), CT: clock time. (**B**) The acute effect of the 1h-light pulse on the total amount of sleep in SA (green) and DCZ (purple) group. The white bar represents the data from the control condition and the gray bar indicates the data from the dark pulse. (**C)** The sleep-wake stage difference between light pulse and control. No difference was found between dark and control condition in both groups. (**D)** and **(E)** Mean spectral distribution of Cortical EEG power density during W, SWS and PS during the period of dark pulse and controls after administering SA and DCZ. To get a better visualization of low frequency spectral power part, the x axis was broken between 35-55 Hz (no significant difference around this frequency). The inserted bars refer to the EEG spectral power band (δ: 0.5-3 Hz, θ:3-9 Hz, spindle: 9-15 Hz, α:15-20 Hz, β:20-30 Hz, γ: 30-60 Hz) during dark and control. Orange represents the controls and black represents the darkness. There is no difference between dark pulse and controls in both groups. Asterisks represent significant differences for post hoc comparisons.

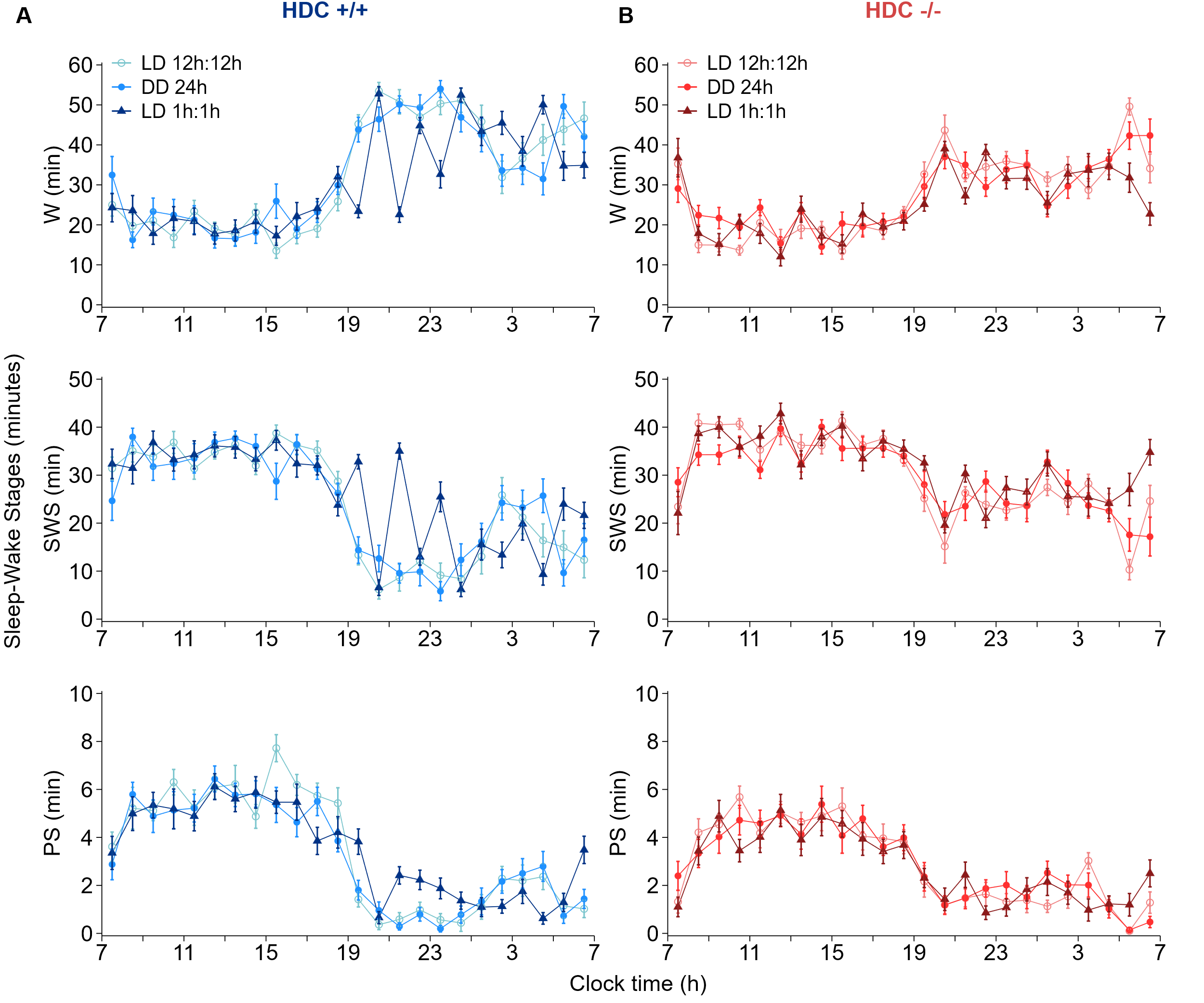

**Figure S2 The sleep-wake stages in HDC+/+ and HDC-/- under three light conditions.**
